# Filling Gaps in Bacterial Amino Acid Biosynthesis Pathways with High-throughput Genetics

**DOI:** 10.1101/192971

**Authors:** Morgan N. Price, Grant M. Zane, Jennifer V. Kuehl, Ryan A. Melnyk, Judy D. Wall, Adam M. Deutschbauer, Adam P. Arkin

## Abstract

For many bacteria with sequenced genomes, we do not understand how they synthesize some amino acids. This makes it challenging to reconstruct their metabolism, and has led to speculation that bacteria might be cross-feeding amino acids. We studied heterotrophic bacteria from 10 different genera that grow without added amino acids even though an automated tool predicts that the bacteria have gaps in their amino acid synthesis pathways. Across these bacteria, there were 11 gaps in their amino acid biosynthesis pathways that we could not fill using current knowledge. Using genome-wide mutant fitness data, we identified novel enzymes that fill 9 of the 11 gaps and hence explain the biosynthesis of methionine, threonine, serine, or histidine by bacteria from six genera. We also found that the sulfate-reducing bacterium *Desulfovibrio vulgaris* synthesizes homocysteine (which is a precursor to methionine) by using DUF39, NIL/ferredoxin, and COG2122 proteins, and that homoserine is not an intermediate in this pathway. Our results suggest that most free-living bacteria can likely make all 20 amino acids and illustrate how high-throughput genetics can uncover previously-unknown amino acid biosynthesis genes.

## Introduction

Although it has been known for decades how the model bacterium *Escherichia coli* synthesizes all 20 of the standard amino acids, novel pathways for amino acid biosynthesis continue to be discovered in other bacteria and in archaea. For example, unlike *E. coli*, most bacteria use tRNA-dependent pathways for the biosynthesis of some of these amino acids (Sheppard et al. 2008). More recent discoveries include a novel pathway for methionine biosynthesis in methanogenic archaea and sulfate-reducing bacteria (Rauch et al. 2014; Kuehl et al. 2014; Allen et al. 2015) and a novel route by which *Pelagibacter ubique*, which is perhaps the most abundant bacterium on earth, synthesizes glycine from a waste product of photosynthetic organisms (Carini et al. 2013).

Our incomplete knowledge of these pathways makes it difficult to accurately predict a bacterium’s minimal growth requirements from its genome sequence. Besides saving experimental effort, such predictions would make it possible to understand the ecological role of bacteria that are difficult to cultivate. For example, (Mee et al. 2014) and (D’Souza et al. 2014) used comparative genomics to predict that most bacteria cannot synthesize all 20 of the standard amino acids. Both groups suggested that most free-living bacteria are reliant on cross-feeding of amino acids that are released by other bacteria. However, neither group tested this experimentally by measuring bacterial growth requirements. In contrast, we recently studied 24 heterotrophic bacteria from 15 different genera (Price et al. 2016) and found that all but one of these bacteria grew in defined minimal media with no amino acids present. If we used the automated method that Mee and colleagues relied on (Chen et al. 2013) and excluded *Escherichia coli,* then we had predictions for bacteria from 12 genera that grow in defined media. All of these bacteria were incorrectly predicted to be auxotrophic for multiple amino acids. In other words, there were many putative gaps in the biosynthetic pathways – enzymatic steps that are required for biosynthesis and were missing from the genome annotation – but these gaps were misleading.

To understand why, we manually examined the predictions for bacteria from 10 different genera. Although the automated method identified a total of 173 gaps, we argue that just 11 of these represent genuine gaps in our biological knowledge. Another gap relates to a recently-discovered and poorly-understood pathway for the biosynthesis of homocysteine (which is a precursor of methionine) in sulfate-reducing bacteria and archaea (Rauch et al. 2014; Kuehl et al. 2014; Allen et al. 2015). To identify the genes that encode these reactions, we used genome-wide mutant fitness assays, in which a pool of ∼40,000-500,000 different transposon mutants is grown together and DNA sequencing is used to measure how each mutant’s abundance changes during growth (Wetmore et al. 2015; Price et al. 2016). Given the genetic data, we looked for genes that were important for fitness in minimal media, but not in rich media, and whose mutants were rescued by the addition of a specific amino acid. Of the 11 genuine gaps, we identified genes to fill nine of them; these novel enzymes explain the biosynthesis of four different amino acids by bacteria from six different genera. And we found that homocysteine synthesis in *Desulfovibrio vulgaris* Miyazaki F required DUF39, NIL/ferredoxin, and COG2122 proteins, as expected from studies of this pathway in other organisms (Rauch et al. 2014; Rauch and Perona 2016; Kuehl et al. 2014). Our genetic data imply that, in contrast to all other known pathways, homoserine is not an intermediate in homocysteine synthesis in *D. vulgaris.*

## Results

### A high rate of error in automated predictions of auxotrophy

(Mee et al. 2014) relied on predicted phenotypes from the Integrated Microbial Genomes web site (https://img.jgi.doe.gov/; (Chen et al. 2013)), while (D’Souza et al. 2014) made their own predictions. We focus our analysis on the IMG predictions because they are publicly available, but the predictions by D’Souza and colleagues have similar issues (see Appendix 1). Among 24 heterotrophic bacteria that grow in defined media without added amino acids and for which we have mutant fitness data, we selected one representative of each genus whose genome is present in IMG. We also excluded the traditional model bacterium *E. coli.* This left us with 12 bacteria, and for each bacterium, IMG incorrectly predicted auxotrophy for at least two amino acids.

We examined 10 of these bacteria in more detail. On average, IMG predicts that these bacteria are prototrophic for 9.6 amino acids, auxotrophic for 6.2 amino acids, and makes no prediction either way for 4.2 amino acids (Figure 1). To verify that these bacteria grow in the absence of any externally provided amino acids, we performed multi-transfer growth experiments for seven of them (Appendix 2). We also tested the vitamin requirements of these seven bacteria (Appendix 2). Six of the bacteria (*Burkholderia phytofirmans* PsJN, *Desulfovibrio vulgaris* Miyazaki F, *Herbaspirillum seropedicae* SmR1, *Marinobacter adhaerens* HP15, *Phaeobacter inhibens* BS107, and *Pseudomonas stutzeri* RCH2) did not require the addition of any amino acids or vitamins for growth. *Sinorhizobium meliloti* 1021 did not grow without added vitamins, which is consistent with a previous report that it requires biotin (but not amino acids) for growth (Watson et al. 2001).

**Figure 1:**
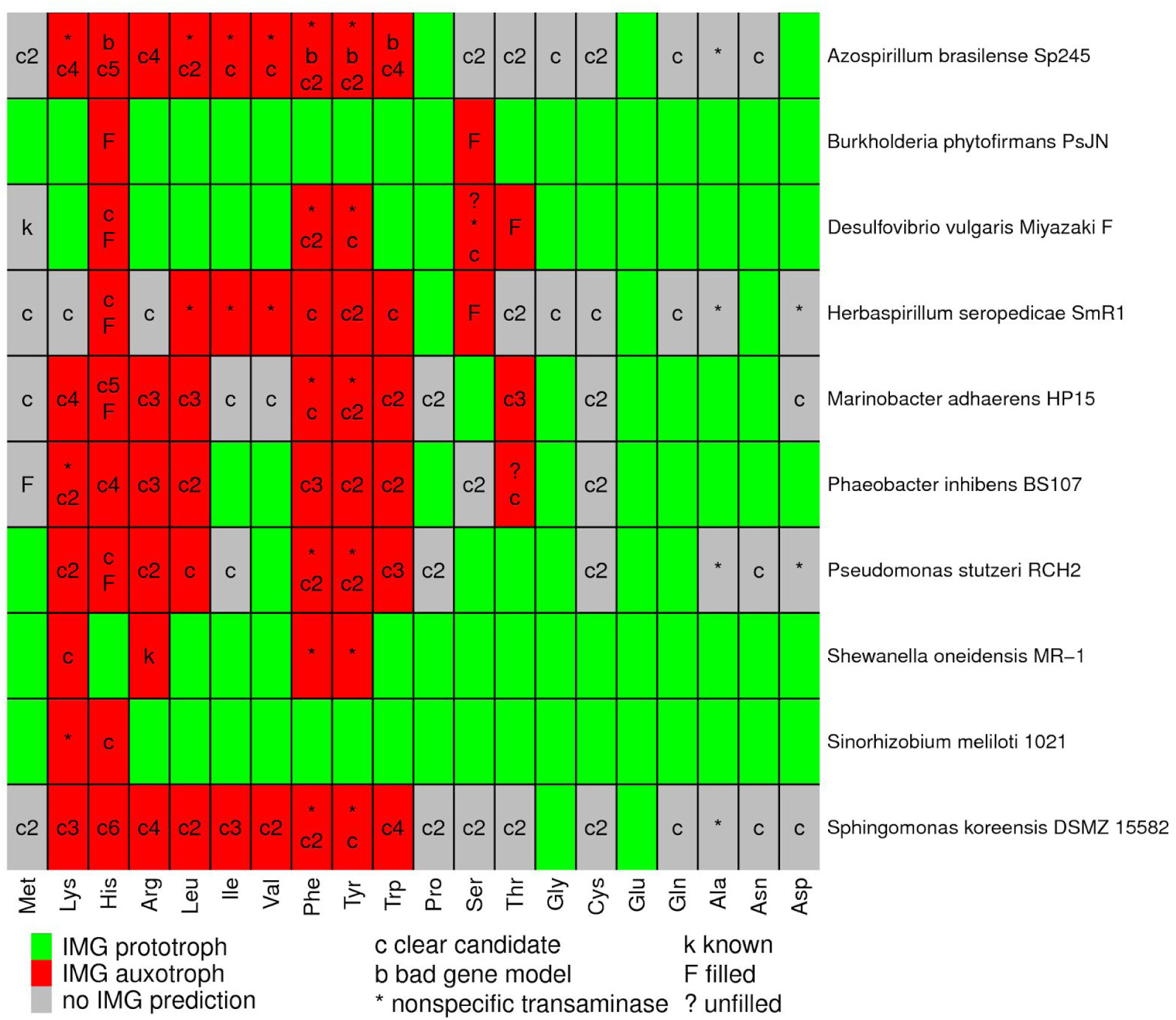
Gaps in amino acid biosynthesis in 10 bacteria. For each amino acid, we identified the missing reactions or gaps in the IMG predictions, and we show a single-letter code with the classification of each gap. A cell may have multiple codes, one for each gap. “Clear candidate” means that at least two annotation resources identified a gene for the gap. If there was more than one gap with a clear candidate, this is shown with a number (i.e., “c2” for two gaps with clear candidates). “Known” means that the step is described in the literature but not in the databases. Some reactions are involved in the biosynthesis of multiple amino acids, so some gaps are shown multiple times.

For each of the 104 cases in which IMG failed to predict that the bacterium could synthesize the amino acid, we used the IMG website to identify the gaps – the reactions that were necessary for biosynthesis of the amino acid but no gene was predicted (in IMG) to encode them. In many cases, a pathway had more than one gap (more than one reaction that is required to make the amino acid was not associated with a gene). Overall, there were a total of 173 gaps in amino acid biosynthesis among the 10 bacteria.

### Of 173 putative gaps in amino acid biosynthesis pathways, 11 are gaps in biological knowledge

140 of the 173 gaps (81%) had clear candidates from other annotation resources: a protein was annotated with the putatively missing enzymatic capability by at least two out of three of TIGRFam (Haft et al. 2013), KEGG (Kanehisa and Goto 2000), and SEED/RAST (Overbeek et al. 2014)). Five of the clear candidates are fused to other biosynthetic enzymes, and these fusions might cause the genes to be missed when searching for bidirectional best hits (Chen et al. 2013). In some other cases, the presence of a potential ortholog is noted on the IMG website, but the gene’s annotation was not deemed high-confidence enough to predict that the pathway is present. These ambiguous cases are labeled as “auxotroph” on the IMG website (as of August 2017) and were so considered in the analyses of (Mee et al. 2014).

To test if these 140 clear candidates were actually involved in amino acid biosynthesis, we examined genome-wide mutant fitness data for each of the 10 bacteria across dozens of growth conditions (Price et al. 2016). In general, a gene that is involved in the biosynthesis of amino acids should be important for fitness in most defined minimal media experiments but not during growth in media that contains yeast extract (which contains all of the standard amino acids) or casamino acids (which contains all of the standard amino acids except tryptophan). For nine of the gaps, more than one clear candidate was identified in the genome and no strong phenotype was found in the mutant fitness data (Supplementary Table 1), which may indicate genetic redundancy. For the remaining 131 steps, we classified 61 genes as auxotrophs because they were important for fitness in most defined media conditions. Some examples are shown in Figure 2. Note that we define gene fitness in a condition as the log_2_ change in the abundance of mutants in that gene after growth from an optical density of 0.02 to saturation (usually 4-8 generations) (Wetmore et al. 2015). Genes that are not important for fitness will have fitness values near zero, and fitness values of under -2 indicate a strong defect in growth. As shown in Figure 2, these auxotrophs had strong fitness defects if amino acids are absent and had little phenotype (fitness near zero) if amino acids were added. Furthermore, in some cases, the gene is expected to be required for the synthesis of just one amino acid, and we measured gene fitness with that amino acid as the sole source of carbon or nitrogen. In these cases, the amino acid rescues the auxotroph, as expected. For example, the bottom right panel in Figure 2 shows that mutants in *hisF* from *Sphingomonas koreensis* (Ga0059261_1048) were rescued when L-histidine was present. Overall, the fitness data shows that these 61 gaps were filled correctly.

**Figure 2:**
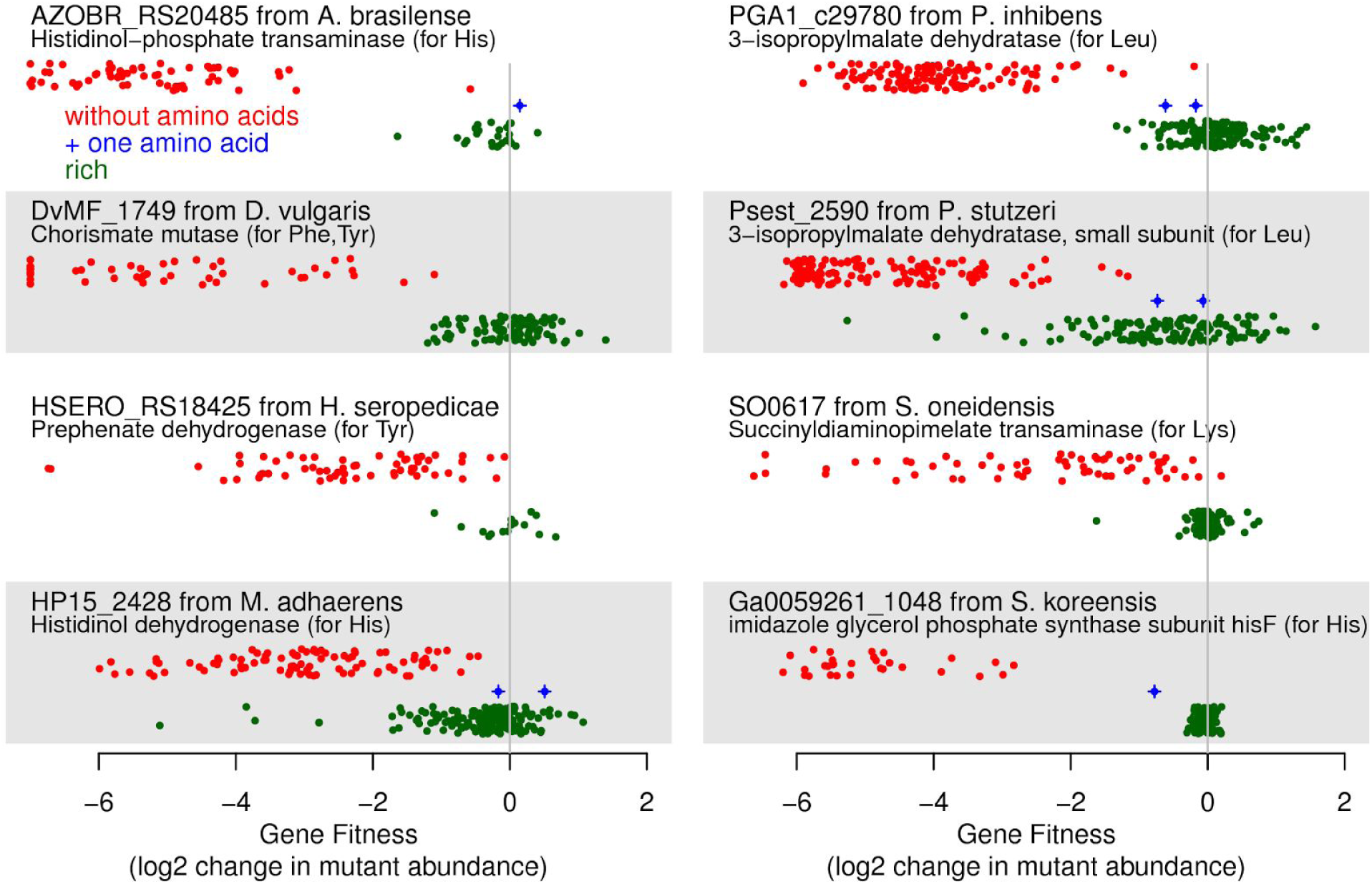
Mutant fitness data for clear candidates that have auxotrophic phenotypes. We selected one gene at random from each organism that has such a gene. In each panel, the *x* axis shows gene fitness (values below -7 are shown at -7), and the *y* axis separates experiments by whether most amino acids were available (green points) or not (red points). In between, we show experiments in which only the relevant amino acid was provided (blue points, if any). Within each category, the *y* axis is random.

Another 54 of the clear candidates were identified as putatively essential for growth in rich media with added amino acids (see Methods). Amino acid biosynthesis genes may be essential in rich media because the bacterium cannot take up the amino acid or because the biosynthetic pathway overlaps with another essential process. For example, in *E. coli*, *dapABDEF* are required for both lysine synthesis and peptidoglycan synthesis and are essential for growth in standard rich media such as LB (Price et al. 2016; Gerdes et al. 2003). In the 10 bacteria under consideration, the clear candidates are much more likely than other proteins to be essential (41% versus 9%), which suggests that most of the essential candidates are truly involved in amino acid biosynthesis.

Among the remaining clear candidate genes, we had insufficient coverage to quantify mutant fitness data for nine (non-essential) genes, and another five may be genetically redundant with other genes (Supplementary Table 1). There were just two clear candidates for which the lack of an auxotrophic phenotype was surprising (Psest_1986 and DvMF_1902), and we identified a potential explanation for Psest_1986 (Appendix 3). Overall, we confirmed 115 of 140 of the clear candidate genes (82%) as being either auxotrophic or essential.

Of the remaining 33 gaps, two gaps were due to an error in the genome sequence or in the identification of a protein-coding gene. First, a missing step in *Azospirillum brasilense* Sp245 is due to a sequencing error that created a frameshift in histidinol dehydrogenase. Nucleotides 1,148,979 to 1,150,284 of the main chromosome (NC_016594.1; (Wisniewski-Dyé et al. 2011)) are very similar (over 80% amino acid identity) to *hisD* (AZL_d03600) from *Azospirillum sp.* B510, but the reading frame is interrupted by a frameshift. This region lies between AZOBR_RS19500 and _RS19510 and is not currently annotated with any genes. Transposon mutants in this region have reduced fitness in defined media (data of (Price et al. 2016)), which suggested that this region of the genome is functional. We sequenced the region on both strands using Sanger sequencing and both reads identified a single nucleotide insertion error at nucleotide 1,149,827 of the published sequence. Once this error is corrected, there is an open reading frame for the complete *hisD* gene (it aligns over its full length and without gaps to AZL_d03600). Second, in the original annotation of *A. brasilense*, which is used in IMG, there is a pseudogene that is annotated as two parts of shikimate kinase (AZOBR_40120, AZOBR_40121). (Shikimate kinase is required for the biosynthesis of aromatic amino acids.) In the updated annotation for the same genome sequence in NCBI’s RefSeq, there is instead a protein-coding gene (AZOBR_RS03225) that is annotated as shikimate kinase. The newly predicted protein does not contain any frameshifts (the error was purely due to gene calling). In a mutant library with 105,000 different transposon insertions in *A. brasilense*, there are no insertions within the 636 nucleotides of AZOBR_RS03225 (Price et al. 2016). This suggests that AZOBR_RS03225 is a genuine and essential protein.

Of the remaining 31 gaps, 18 were transaminase reactions, which may be nonspecific (Lal et al. 2014). For example, IMG lists aromatic amino-acid transaminase as the necessary gene for the final step in the biosynthesis of phenylalanine and tyrosine, but *E. coli* contains multiple transaminases with overlapping substrate specificities that can perform these steps. *E. coli tyrB* and *aspC* contribute to the synthesis of tyrosine and phenylalanine; *ilvE* contributes to phenylalanine synthesis; and all three of these genes contribute to the synthesis of other amino acids as well (Pittard and Yang, 2008). IMG predicted that aromatic amino-acid transaminase is missing in eight of the ten bacteria we examined, but all eight of these bacteria contain multiple amino acid transaminase genes. As another example, IMG lists N-succinyldiaminopimelate aminotransferase as required for the synthesis of lysine via succinylated intermediates. In *E. coli,* this activity is provided by both *argD* and *serC*, which also catalyze transamination reactions in the biosynthesis of other amino acids (Lal et al. 2014) (also reviewed in EcoCyc, (Keseler et al. 2005)). Similarly, in *Azospirillum brasilense* Sp245 and in *Phaeobacter inhibens* BS107, a putative ornithine transaminase (*argD*; AZOBR_RS19025 or PGA1_c24230) may provide the missing N-succinyldiaminopimelate aminotransferase activity. This gene is essential in both organisms (Price et al. 2016), which is consistent with our proposal because this step is also required for peptidoglycan synthesis. Because many amino acid transaminases are non-specific and because their specificity is currently difficult to annotate, the absence of a transaminase should not be used to predict auxotrophy.

After removing the transaminase reactions, 13 gaps remained, but two of these gaps had already been filled by experimental studies. First, in *Shewanella oneidensis* MR-1, SO3749 is the missing acetylornithine deacetylase for arginine synthesis (Deutschbauer et al. 2011). Second, the sulfate-reducing bacterium *Desulfovibrio vulgaris* Miyazaki F contains recently-discovered genes for the biosynthesis of homocysteine, which is a precursor to methionine (DUF39, NIL/ferredoxin, and COG2122; (Rauch et al. 2014) http://f1000.com/work/citation?ids=2169833&pre&suf&sa=0 (Kuehl et al. 2014) (Rauch and Perona 2016)). Unfortunately, the information from these studies has not made its way into the annotation databases.

Of the 173 gaps in amino acid biosynthesis from the automated tool, just 11 represented genuine gaps in biological knowledge. For 9 of these 11 gaps, we provide genetic evidence for the genes that provide the missing enzymatic capabilities. In addition, our genetics data for *Desulfovibrio vulgaris* Miyazaki F provides additional insights into the recently-discovered pathway for methionine. We will first describe methionine synthesis in *D. vulgaris* in more detail and then each of the 9 gaps that we filled using high-throughput genetics data.

### Methionine synthesis in Desulfovibrio vulgaris

We recently found that a DUF39 protein (DUF is short for domain of unknown function) is required for homocysteine formation in *Desulfovibrio alaskensis* (Kuehl et al. 2014). A genetic study in the methanogen *Methanosarcina acetivorans* (Rauch et al. 2014) also found that a DUF39 protein (MA1821) is involved in homocysteine synthesis, along with a protein containing NIL and ferredoxin domains (MA1822). (The NIL domain is named after a conserved subsequence and the PFam curators suggest that it might be a substrate binding domain.) Although the molecular function of these proteins is unclear, biochemical studies of cell extracts suggest that in methanogens, homocysteine is formed by reductive sulfur transfer to aspartate semialdehyde, and this process requires the DUF39 and/or the NIL/ferredoxin proteins (Allen et al. 2015). In contrast, in the well-characterized pathways for methionine synthesis, aspartate semialdehyde is converted to homocysteine via multiple steps: aspartate semialdehyde is first reduced to homoserine by homoserine dehydrogenase; then the alcohol group is activated by acetylation, succinylation, or phosphorylation; and finally the sulfide is transferred to form homocysteine.

The genome of *Desulfovibrio vulgaris* Miyazaki encodes DUF39 and NIL/ferredoxin proteins and does not appear to encode any of the well-characterized pathways for methionine biosynthesis, so we expected that it would synthesize methionine by the recently discovered pathway. Indeed, we found that the DUF39 protein (DvMF_1464) and the NIL/ferredoxin protein (DvMF_0262) were important for growth in minimal media and that mutants in these genes were rescued by added methionine (Figure 3). As far as we know, this is the first experimental evidence that the NIL/ferredoxin protein is required for methionine synthesis in *Desulfovibrio*.

**Figure 3:**
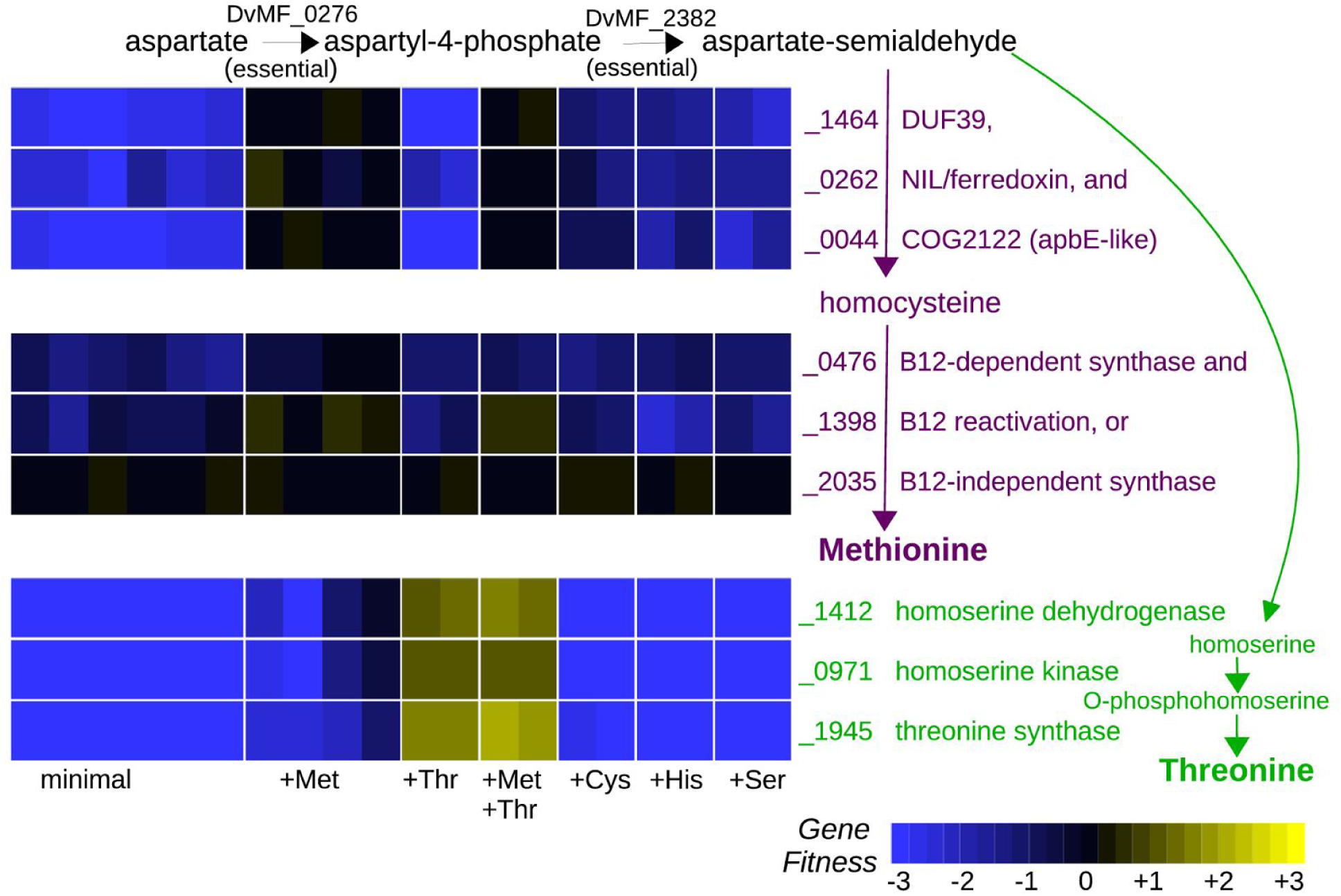
Synthesis of methionine and threonine in Desulfovibrio vulgaris Miyazaki F. We grew a pool of transposon mutants of *D. vulgaris* in a defined lactate/sulfate medium with or without added amino acids. L-methionine or L-threonine were supplemented at either 1 mM and 10 mM. D,L-cysteine, D,L-histidine, or L-serine were supplemented at 1 mM.

We also found that homoserine dehydrogenase (DvMF_1412), which is required for homoserine synthesis, was important for fitness in minimal media (Figure 3). This was expected because homoserine is an intermediate in the biosynthesis of threonine as well as in the standard pathway of homocysteine biosynthesis. Furthermore, supplementing the medium with threonine alone was sufficient to rescue the homoserine dehydrogenase mutants (Figure 3), which implies that homoserine dehydrogenase is not required for methionine synthesis. This confirms that homoserine is not an intermediate in the DUF39 pathway for homocysteine synthesis, as previously suggested based on a biochemical study of cell extracts from methanogens (Allen et al. 2015).

Comparative genomics analyses had also suggested that COG2122 might be involved in the this pathway (Kuehl et al. 2014) (Rauch et al. 2014). (COG is short for clusters of orthologous groups and COG2122 is distantly related to ApbE proteins (Galperin et al. 2015).) We found that the COG2122 protein in *D. vulgaris* (DvMF_0044) was important for growth in defined media and that mutants were rescued by added methionine (Figure 3). We also found that, across a variety of growth conditions, the fitness of the COG2122 protein was virtually identical to that of the DUF39 protein (r = 0.98 across 170 fitness experiments). This strongly suggests that the COG2122 protein is also required for homocysteine formation. In contrast, (Rauch and Perona 2016) found that the orthologous protein from *Methanosarcina acetivorans (*MA1715) was important for growth in defined media at low sulfide concentrations, but that mutants could be rescued by either higher sulfide concentrations or by added cysteine or homocysteine. They proposed that COG2122 aids in the formation of an unknown intermediate in sulfide assimilation for both methionine and cysteine. Because we grew *D. vulgaris* with sulfate as the electron acceptor (which is reduced to sulfide), and because we added 1 mM sulfide to the media as a reductant, it seemed implausible that COG2122 would be important in *D. vulgaris* because of low sulfide concentrations. Nevertheless, we performed a fitness experiment with added D,L-cysteine. We found that in *D. vulgaris*, mutants of all three homocysteine biosynthesis genes were partially rescued by added cysteine (Figure 3; mean fitness = -1.0). This suggests that *D. vulgaris* might have a minor alternate route to homocysteine, perhaps via a putative cystathionine β-lyase (DvMF_1822). In any case, we propose that COG2122 is required for the conversion of aspartate semi-aldehyde and sulfide to homocysteine; its apparent dispensability for homocysteine formation in *M. acetivorans* under some conditions might be due to genetic redundancy.

In summary, we identified three genes in *D. vulgaris* that are required for homocysteine synthesis, as expected from previous studies of another species of *Desulfovibrio* and of methanogens. We provided genetic evidence that homoserine is not an intermediate in this pathway and that COG2122 is required.

### A homoserine kinase for threonine synthesis in Desulfovibrio vulgaris that is related to shikimate kinase

IMG predicts that *D. vulgaris* is a threonine auxotroph because of a missing homoserine kinase. We identified mutants in three genes as being rescued by added threonine: threonine synthase (DvMF_1945), homoserine dehydrogenase (DvMF_1412), and DvMF_0971, which was originally annotated as a shikimate kinase (Figure 3). This observation suggests that DvMF_0971 might instead be a homoserine kinase, as both reactions involve the phosphorylation of an alcohol group. Indeed, the genome encodes another shikimate kinase (DvMF_1410) which appears to be essential, as are other genes in the chorismate synthesis pathway (DvMF_1750, DvMF_0373, DvMF_0962, DvMF_1748, and DvMF_1408). Furthermore, DvMF_1410 is similar to the shikimate kinase II from *E. coli* (47% identical), while DvMF_0971 is more distantly related (27% identical). To test our prediction that DvMF_0971 is a homoserine kinase, we cloned it and transformed it into a *thrB*- strain of *E. coli* from the Keio deletion collection (Baba et al. 2006). This strain of *E. coli* does not grow in minimal media due to a lack of homoserine kinase activity, and its growth was rescued by the expression of DvMF_0971. Thus, DvMF_0971 is the homoserine kinase of *D. vulgaris*.

### A split methionine synthase in Phaeobacter inhibens

The IMG website makes no prediction as to whether *Phaeobacter inhibens* DSM 17395 (formerly *P. gallaeciensis*; also known as strain BS107) can synthesize methionine because it was unable to identify a gene for methionine synthase. We propose that *P. inhibens* contains a vitamin B12-dependent methionine synthase that is split into three genes (Figure 4A). For comparison, in *E. coli*, the vitamin B12-dependent methionine synthase (MetH) contains a methyltransferase domain (PF02574), a pterin binding domain (PF00809), a vitamin B12 binding cap (PF02607), and a vitamin B12 binding domain (PF02310). In *P. inhibens,* PGA1_c13370 has the methyltransferase domain, PGA1_c16040 has the pterin-binding domain, and MtbC (PGA1_c13350) was originally annotated as “putative dimethylamine corrinoid protein” and contains the vitamin B12-binding cap and vitamin B12-binding domains. (Thole et al. 2012) previously suggested that two of these proteins might be involved in methionine synthesis. We found that all three of these genes were important for growth in defined media, and their mutants were rescued by the addition of methionine or of casamino acids (Figure 4B). Furthermore, these genes had similar phenotypes across 270 diverse fitness experiments (all r > 0.7, Pearson correlations). This confirms that they work together to provide the missing methionine synthase activity.

**Figure 4:**
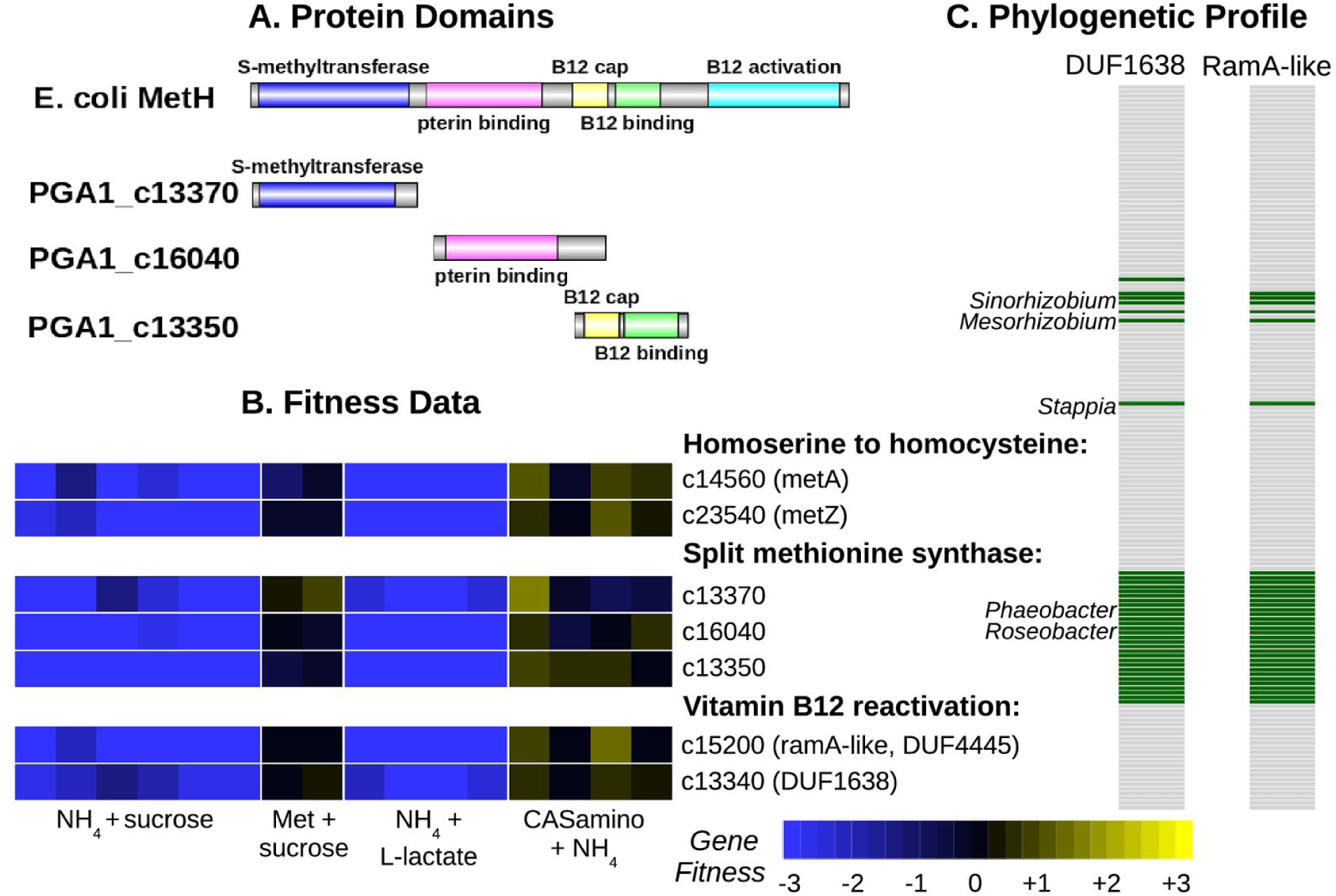
Methionine synthesis in *Phaeobacter inhibens* by a three-part methionine synthase and two vitamin B12 reactivation proteins. (A) Domain content of MetH from *E. coli* and of the three-part methionine synthase of *P. inhibens.* (B) Fitness data of the methionine synthesis genes. We grew pools of mutants of *P. inhibens* aerobically in defined media with a variety of carbon and nitrogen sources. (C) Phylogenetic profile of the presence or absence of the vitamin B12 reactivation proteins across 158 α-Proteobacterial genomes from MicrobesOnline (Dehal et al. 2010). The bacteria are ordered by evolutionary relationships and some of the genera that contain these proteins are labeled.

### A RamA-like protein for vitamin B12 activation in Phaeobacter and Desulfovibrio

The activity of vitamin B12-dependent methionine synthase also requires the “reactivation” of vitamin B12 to reduce co(II)balamin, which can form as a side reaction of this enzyme, to co(I)balamin. In *E. coli*, the reactivation of B12 is provided by yet another domain (PF02965) at the C terminus of the MetH protein, but in other bacteria this can be a separate protein. However no member of PF02965 was found in *P. inhibens* or in related bacteria such as *Dinoroseobacter shibae*. (Thole et al. 2012) proposed that PGA1_c13360, which contains a radical SAM domain, might be involved in B12 activation, but we found that this gene was not important for growth in minimal media (all fitness values were within -0.5 to +0.5). Instead, we identified two other genes that had correlated fitness with the other methionine synthase genes and are likely to be involved in B12 reactivation: a protein with ferredoxin and DUF4445 domains (PGA1_c15200) and a DUF1638 protein (PGA1_c13340). As shown in Figure 4B, mutants in these genes are rescued by added methionine.

The DUF4445 protein is distantly related to RamA, which uses ATP to drive the reductive activation of corrinoids in methanogens (Ferguson et al. 2009). Indeed, Ferguson and colleagues predicted that bacterial homologs of RamA would be involved in vitamin B12 reactivation, and we previously proposed that in *Desulfovibrio alaskensis*, a RamA-like protein (Dde_2711) would be involved in B12 reactivation because it is cofit with MetH (r = 0.90; (Kuehl et al. 2014)). We also found evidence that DUF4445 is involved in the reactivation of B12 in the *D. vulgaris*, which contains a B12-dependent methionine synthase (DvMF_0476) that lacks the standard B12 activation domain. This methionine synthase has a very similar fitness pattern as DvMF_1398, which contains two DUF4445 domains (r = 0.92 across 170 experiments; also see Figure 3). We infer that DUF4445 proteins perform the reactivation of vitamin B12 in diverse bacteria.

We do not have a specific proposal for the function of the DUF1638 protein. Although it is adjacent to the gene that encodes the vitamin B12-binding cap and vitamin B12-binding domains, the DUF1638 protein is downstream and at the end of the operon, so its phenotype is unlikely to be due to polar effects. We thought that DUF1638 could be involved in the synthesis of vitamin B12 rather than in B12 reactivation *per se*, but unlike the DUF1638 protein, the genes in the vitamin B12 synthesis pathway have few insertions and are probably essential in *P. inhibens.* (The CobIGJMKFLHBNSTQDPV proteins are all essential, as is one of two CobO-like proteins. Vitamin B12 synthesis may be essential, even when methionine is provided, because of a vitamin B12-dependent ribonucleotide reductase.) Across the α-Proteobacteria, the presence or absence of DUF1638 is nearly identical to that of the RamA-like protein (Figure 4C), which is consistent with a close functional relationship. One unexplained aspect of this distribution is that some of the bacteria with the RamA-like and DUF1638 proteins also contain a MetH that includes a standard reactivation domain (i.e., *Sinorhizobium meliloti* 1021). Overall, we identified five genes that are involved in methionine synthesis in *P. inhibens*: three pieces of methionine synthase and two proteins for the reactivation of vitamin B12, including a RamA-like protein that is also involved in vitamin B12 reactivation in *Desulfovibrio*.

### An “FAD-linked oxidase” is involved in serine synthesis

In *Burkholderia phytofirmans* and in *Herbaspirillum seropedicae*, we were unable to find the standard D-phosphoglycerate dehydrogenase (*serA*), which catalyzes the first step in serine biosynthesis. We propose that another oxidase provides this missing activity: BPHYT_RS03150 in *B. phytofirmans* or HSERO_RS19500 in *H. seropedicae.* These two proteins are very similar (76% amino acid identity) and both were originally annotated as “FAD-linked oxidase.” They contain an N-terminal DUF3683 domain, FAD-binding and FAD oxidase domains, a 4Fe-4S dicluster domain, two cysteine-rich CCG domains (which are often associated with redox proteins), and a C-terminal DUF3400 domain.

In *B. phytofirmans*, this oxidase was important for fitness in most defined media, but not in rich media (LB), or when casamino acids were added, or when L-serine was the nitrogen source (Figure 5A). If we supplemented our standard glucose/ammonia minimal media with L-serine, then mutants in this gene were partially rescued, as were mutants in the gene for the next step in serine synthesis (phosphoserine transaminase or *serC*; Figure 5B). *B. phytofirmans* may preferentially uptake ammonia instead of serine, which could explain why the two serine synthesis genes are only partially rescued by added serine if ammonia is present. The gene for the final step in serine synthesis (phosphoserine phosphatase or *serC*, BPHYT_RS09200) may be important for fitness even in rich media, as mutant strains were at low abundance in our pool of mutants.

**Figure 5:**
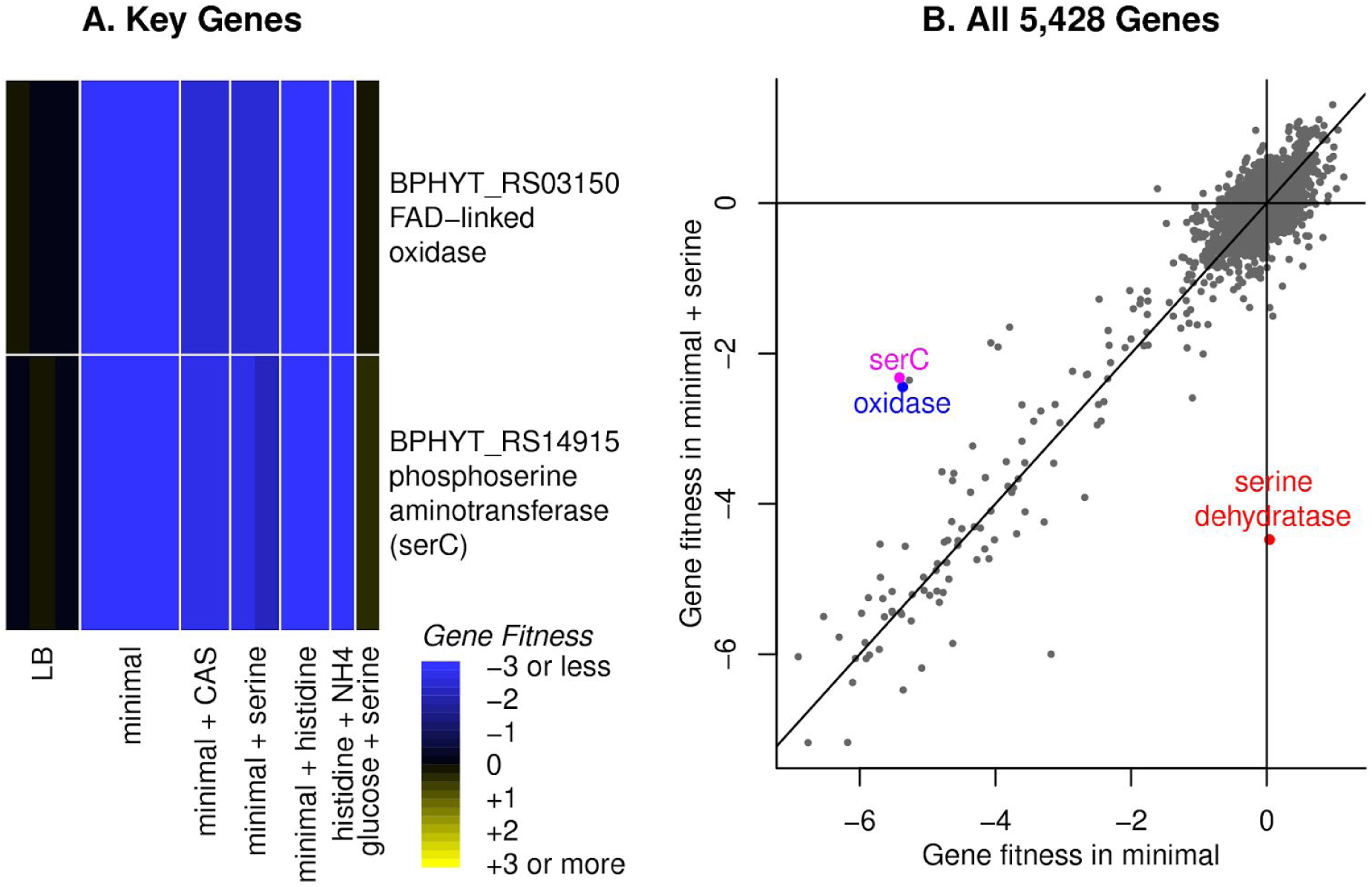
Serine biosynthesis in *Burkholderia phytofirmans*. **(A)** A heatmap of gene fitness of the putative phosphoglycerate dehydrogenase (“FAD-linked oxidase”) and another serine biosynthesis gene (the aminotransferase *serC*). Our standard minimal media for this organism contains glucose and ammonia, and CAS is short for casamino acids, which contains both L-serine and L-histidine. **(B)** A comparison of gene fitness during growth in minimal media (*x* axis) or in media that was supplemented with 1 mM L-serine (*y* axis). The lines show *x* = *0*, *y* = *0*, or *x* = *y*. We highlight the genes from part (A) as well as the catabolic serine dehydratase. The unmarked point near *serC* and the dehydrogenase is a putative cell wall synthesis gene (BPHYT_RS14855, ADP-L-glycero-D-manno-heptose-6-epimerase). Each point shows the average of two replicate experiments.

In *H. seropedicae,* this oxidase appears to be essential in rich media with amino acids. Mutants in the phosphoserine transaminase (HSERO_RS18435) and the phosphoserine phosphatase (HSERO_RS03150) were also at low abundance in our mutant pool, so the poor viability of mutants in HSERO_RS19500 is consistent with a role in serine synthesis. To look for other candidates for this step, we collected fitness data for *H. seropedicae* in minimal media with and without added L-serine, but we did not identify any genes whose mutants were rescued by added L-serine. (Averaging across two replicate experiments, there were no genes with fitness under -2 in minimal glucose media and fitness above -1 in minimal glucose media that was supplemented with 1 mM L-serine.)

We predict that in both bacteria, the phosphoglycerate dehydrogenase activity is provided by the FAD-linked oxidase that has additional DUF3683, CCG, and DUF3400 domains. However, neither BPHYT_RS03150 nor HSERO_RS19500 complemented the growth deficiency of a *serA*-strain of *E. coli* from the Keio collection in minimal media (Baba et al. 2006). The FAD-linked oxidase might require another cofactor or protein for activity, or it might have some other unexpected role in serine synthesis. Both organisms contain genes for both of the other steps in serine synthesis (the phosphoserine transaminase *serC* and the phosphoserine phosphatase *serB*), and these genes are either essential or their mutants are auxotrophic, so we do not expect the FAD-linked oxidase to be involved in these other steps.

### Histidinol-phosphate phosphatases for histidine synthesis that are similar to phosphoserine phosphatases

We propose that in four of the 10 bacteria, genes that were originally annotated as phosphoserine phosphatases provide the missing histidinol-phosphate phosphatase activities. Phosphoserine phosphate and histidinol phosphate are both of the form R-C(NH_3_ ^+^)-CH_2_OPO_3_ ^2-^, so this is biochemically plausible. These genes are: BPHYT_RS03625 from *Burkholderia phytofirmans*; Psest_3864 from *Pseudomonas stutzeri* RCH2; HP15_461 from *Marinobacter adhaerens* HP15; and HSERO_RS03150 from *Herbaspirillum seropedicae* SmR1. In three of the four bacteria, this gene was important for fitness in minimal media but not in minimal media that was supplemented with histidine (Figure 6A, 6B, 6C). In *H. seropedicae,* mutants in HSERO_RS03150 are at low abundance in our pools, so we do not have fitness data for it. The genes in *H. seropedicae* whose mutants were rescued by added histidine are all annotated as performing other steps in histidine biosynthesis (Figure 6D), so our data does not suggest another candidate for this step. The four putative histidinol-phosphate phosphatases are all similar to each other (≥45% amino acid identity), so the results from the various bacteria corroborate each other.

**Figure 6:**
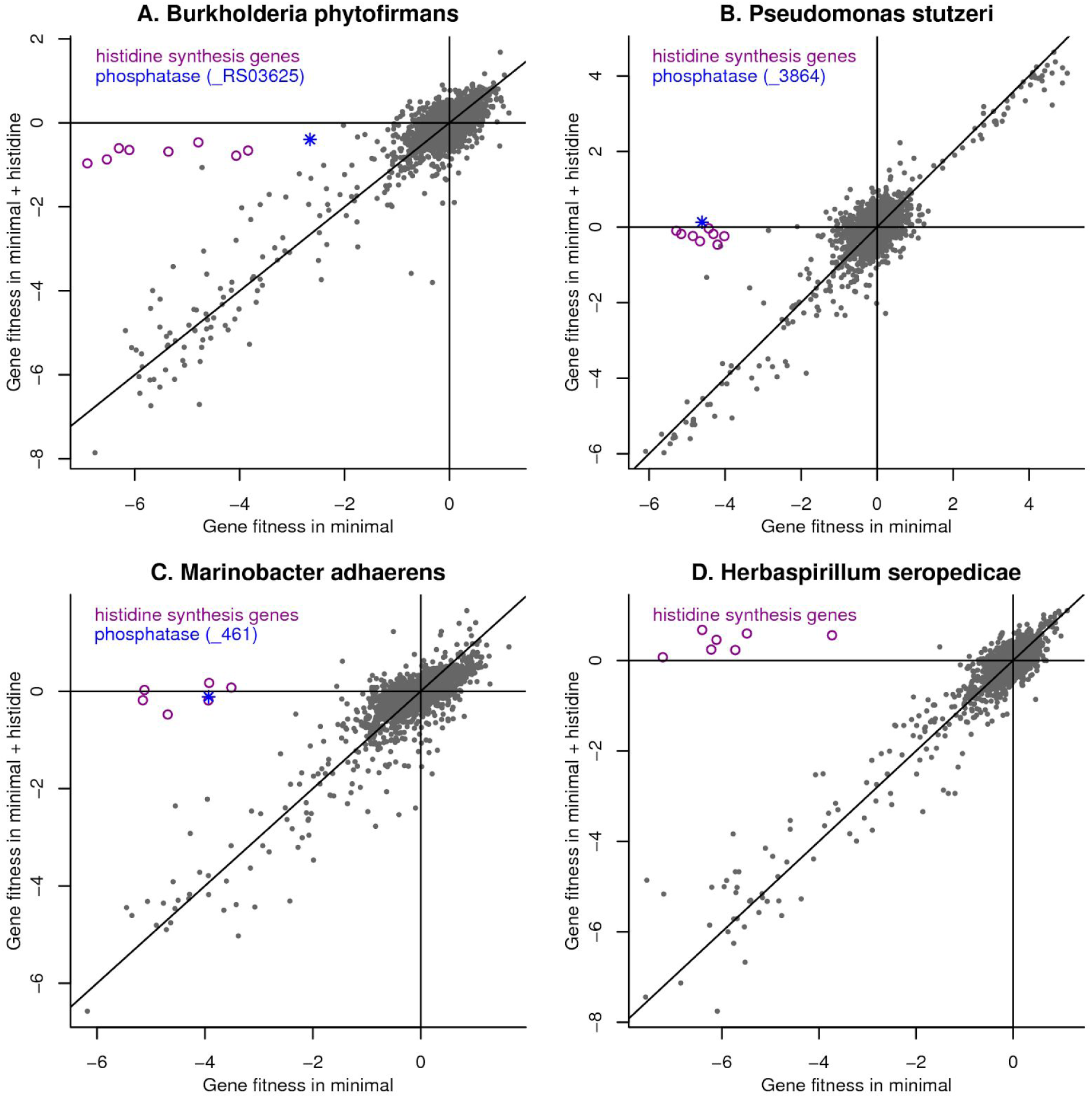
Identification of novel histidinol-phosphate phosphatases. We compared gene fitness in minimal media (*x* axis) or in minimal media supplemented with 1 mM L-histidine (*y* axis) for (A) *Burkholderia phytofirmans*, (B) *Pseudomonas stutzeri*, (C) *Marinobacter adhaerens*, and (D) *Herbaspirillum seropedicae*. In each panel, we highlight the putative phosphatase as well as genes for other steps in histidine biosynthesis. For *H. seropedicae*, the phosphatase is not shown because of a lack of data. The lines show *x*=*0*, *y*=*0*, and *x*=*y*. Each point shows the average gene fitness from two replicate experiments.

We also note that each of these bacteria contain another gene that is annotated as phosphoserine phosphatase (BPHYT_RS09200, Psest_0489, HP15_2518, and HSERO_RS15175). We believe that these genes are correctly annotated, but we only have fitness data for one of them. We found that Psest_0489 from *P. stutzeri* is important for fitness in most but not all defined media conditions. It is possible that another enzyme in this organism also has phosphoserine phosphatase activity: Psest_2327 is 89% identical to ThrH from *P. aeruginosa*, which has this activity (Singh et al. 2004). If so, this would explain why Psest_0489 is not important for fitness in some defined media conditions with no serine added.

### A phosphoribosyl-ATP diphosphatase from the MazG family

The mutant fitness data for *D. vulgaris* did not identify a candidate for phosphoribosyl-ATP diphosphatase, which is required for histidine biosynthesis. By sequence analysis, we identified DvMF_3078 as a candidate, but we did not have fitness data for this gene. DvMF_3078 is related to MazG (nucleotide pyrophosphatase), which performs a similar reaction. (In both reactions, a nucleotide 5’-triphosphate is converted to a nucleotide 5’-monophosphate.) The ortholog of DvMF_3078 in *D. alaskensis* G20 (Dde_2453) is important for fitness in minimal media and its fitness pattern is most similar to that of *hisI* (r = 0.96; data of (Kuehl et al. 2014), (Price et al. 2014)). These observations suggested that DvMF_3078 encodes the missing phosphoribosyl-ATP diphosphatase.

To test this hypothesis, we studied a transposon mutant of the orthologous gene from *D. vulgaris* Hildenborough (DVU1186). We found that a transposon mutant of DVU1186 (strain GZ8414) grew little if at all in minimal media and that its growth was rescued by the addition of 0.1 mM L-histidine (Figure 7). As a control, we also tested a transposon mutant in DVU2938 (a DUF39 protein which is involved in methionine synthesis) and found that it was unable to grow in defined media even if histidine was added (Figure 7). Thus, in the genus *Desulfovibrio,* a MazG family protein is required for histidine biosynthesis and is probably the missing gene for phosphoribosyl-ATP diphosphatase. It is interesting to note that the MazG family is distantly related to HisE, which provides the phosphoribosyl-ATP diphosphatase activity in most bacteria (Moroz et al. 2005).

**Figure 7:**
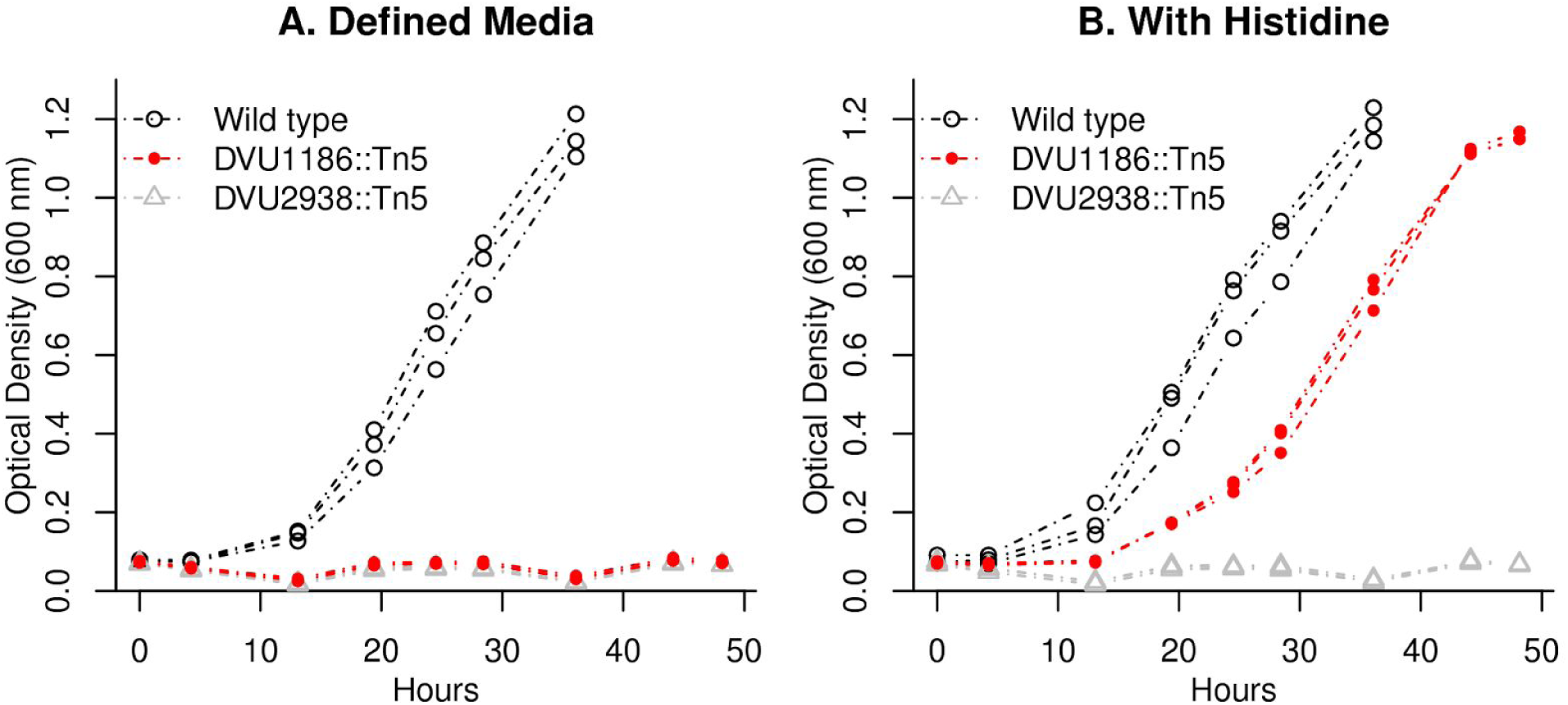
DVU1186 is required for histidine synthesis. We grew transposon mutants of DVU1186 (strain GZ8414) and of DVU2938 (strain GZ9865), as well as wild type *D. vulgaris* Hildenborough, in defined lactate-sulfate medium (panel A) or in the same medium with 0.1 mM L-histidine added (panel B). For each strain and condition, we show three replicates.

### Two unfilled gaps

Two of the 11 genuine gaps remain unfilled. First, we did not identify the phosphoserine phosphatase in *D. vulgaris*. We thought that this activity might be provided by DvMF_0940, which is annotated as histidinol-phosphate phosphatase and could be bifunctional. We also thought that a putative phosphatase (DvMF_1903) that lies downstream of the phosphoglycerate dehydrogenase (which is the previous step in serine synthesis) was a plausible candidate. Unfortunately, we do not have fitness data for either of these genes. We cloned each of these proteins into an *E. coli serB*-strain from the Keio collection, which lacks phosphoserine phosphatase, and neither of the proteins from *D. vulgaris* were able to rescue its growth in minimal media. As another test, we constructed a deletion of DVU0338 from *D. vulgaris* Hildenborough, which is similar to the putative phosphatase DvMF_1903 (77% amino acid identity). We found that the DVU0338-strain could grow in defined media. We suspect that some other protein provides the phosphoserine phosphatase activity in *D. vulgaris.*

Second, the fitness data did not identify a candidate for homoserine kinase in *Phaeobacter inhibens*. We thought that the shikimate kinase (PGA1_c14090), which is essential, might be bifunctional and act on homoserine as well. However, when we cloned this protein into an *E. coli thrB-* strain from the Keio collection, which lacks homoserine kinase, it did not rescue growth in minimal media.

## Discussion

### Many gaps in our understanding of amino acid biosynthesis

Although amino acid biosynthesis is well understood in model organisms such as *Escherichia coli*, our results imply that there are many steps that remain unknown, even in the relatively well-studied Proteobacteria. Once genome-wide fitness data from more diverse bacteria is available, we hope to explain many more mysteries. For example, at least one more pathway of homocysteine synthesis remains to be discovered: thermophilic autotrophs from several divisions of bacteria (i.e., *Aquifex aeolicus*, *Pyrolobus fumarii* 1A, and *Acidimicrobium ferrooxidans* DSM 10331) contain neither the traditional nor the DUF39 pathways of homocysteine biosynthesis, nor the protein thiocarboxylate pathway (Krishnamoorthy and Begley 2011).

Given our limited knowledge, it may be premature to try to predict whether bacteria can synthesize amino acids from their genome sequences. We did identify some recurring types of gaps that should not be used to predict auxotrophy. If we exclude the gaps from IMG that had clear candidates from other annotation resources or were due to errors in gene models, then there were 31 gaps, of which 18 were transaminases, 5 were phosphatases, and 3 were kinases. We cannot determine if this information would lead to better predictions of auxotrophies, as we only studied prototrophic bacteria. If the growth requirements were known for large numbers of bacteria along with their genome sequences, it should become possible to make useful predictions.

### Most free-living bacteria can synthesize all 20 amino acids

The heterotrophic bacteria that we studied are not a random sample of all heterotrophic bacteria. However, of the heterotrophic bacteria that we studied previously, 23 of 24 grew in minimal media (Price et al. 2016), and we selected these on the basis of their genetic tractability rather than their growth requirements. Furthermore, as far as we know, these bacteria were isolated and propagated in media that was supplemented with yeast extract, such as LB, R2A, or marine broth. We do not see why these media would select for prototrophic bacteria. Our tentative conclusion is that most free-living bacteria can synthesize all 20 amino acids, even though we do not know how.

We propose that even free-living bacteria with reduced genomes can synthesize all 20 amino acids. For example, consider the abundant ocean bacterium *Pelagibacter ubique*, which has a streamlined genome and has just 1,354 protein-coding genes (Giovannoni et al. 2005). *P. ubique* has unusual nutritional requirements for reduced sulfur compounds and for glycolate, but given these compounds, it can make all 20 amino acids (Carini et al. 2013). These compounds are released by photosynthetic organisms in the ocean, so it appears that *P. ubique* synthesizes all 20 amino acids in nature. In contrast, our impression is that auxotrophies are widespread in dedicated pathogens and in endosymbionts.

Under the black queen hypothesis, dependencies between organisms can be selected for if the capability is “leaky” and benefits other organisms nearby (Morris et al. 2012). For example, an organism that degrades a toxic compound will also reduce the concentration of that compound that is experienced by its neighbors. It has been suggested that this mechanism could favor the loss of amino acid synthesis genes (D’Souza et al. 2014), but we argue that amino acid synthesis is not so leaky. Even if small amounts of amino acid or protein leak out of nearby cells, it seems questionable that this would provide adequate amino acids for growth, given that about half of the dry weight of bacteria is protein (BioNumbers 101955; (Milo et al. 2010)). And although some mutant strains of *E. coli* will secrete amino acids in sufficient quantities to maintain the growth of auxotrophic strains (Pande et al. 2014), we do not know of any evidence that this is occurs in nature, except for endosymbionts.

Although we are skeptical about the idea that bacteria cross-feed each other amino acids, there is some evidence for the cross feeding of vitamins (Seth and Taga 2014). Because bacteria need vitamins at far lower concentrations than they need amino acids, it seems more plausible that the black queen mechanism could apply to vitamins. Alternatively, because vitamins are present at low concentrations, they might be preferentially recycled from lysed cells rather than broken down for energy. If a subset of bacteria synthesize vitamins rather than taking them up and release them when they die, then many other bacteria would not need to synthesize vitamins. (Even if vitamins are available, some bacteria might be selected to synthesize them if they require relatively high amounts of the vitamin for their metabolism or if vitamin receptors are targeted by phage.) Nevertheless, of the seven bacteria we tested, six grew without added vitamins. Most free-living bacteria may not require exogenous vitamins for growth either.

## Materials and Methods

### Comparison to IMG predictions

We began with 24 heterotrophic bacteria from 15 genera that we had collected large-scale mutant fitness data for (Price et al. 2016). We had previously found that 23 of these bacteria grew in defined media without added amino acids (Price et al. 2016). Since that study was conducted, we also generated a mutant library in *Herbaspirillum seropedicae* SmR1 (see below), which is a plant-associated (endophytic) and nitrogen-fixing bacterium. Although all of these bacteria have been sequenced, not all of them are available in the IMG website, so not all of them have auxotrophy predictions. Also, we arbitrarily selected one representative of each genus. This left us with 13 bacteria. One of these is *Escherichia coli*, which is a traditional model organism and, not surprisingly, IMG correctly predicts that it can make all 20 amino acids. Also, we did not do a detailed analysis for *Dinoroseobacter shibae* DFL-12 or *Dechlorosoma suillum* PS (also known as *Azospira suillum* PS). These had 5 and 6 spurious auxotrophies, respectively, which is similar to numbers for the other bacteria that we did a detailed analysis of. Predictions of amino acid synthesis capabilities were taken from the IMG web site (https://img.jgi.doe.gov/) on May 20, 2016.

### Strains and Media

Except for *H. seropedicae*, mutant libraries were described in (Price et al. 2016). The original sources of the strains are given in Table 1 along with the standard minimal media that we used for each organism and the standard carbon source that we used. These media all contain ammonium chloride as the standard nitrogen source, but this was omitted for some nitrogen source experiments. The minimal media also contain inorganic salts, buffer, and either Wolfe’s vitamins or Thauer’s vitamins. Media components are given in the supplementary material of (Price et al. 2016) and are available for each experiment in the Fitness Browser (http://fit.genomics.lbl.gov/).

**Table 1:** Sources of wild-type strains and their minimal media.

| Strain | Source | Minimal media |
| --- | --- | --- |
| *Azospirillum brasilense* Sp245 | Stephen Fairclough, JGI | RCH2_defined_noCarbon + glucose |
| *Burkholderia phytofirmans* PsJN | DSM 17436 | RCH2_defined_noCarbon + glucose |
| *Desulfovibrio vulgaris* Miyazaki F | Terry Hazen, ORNL | MoLS4_no_lactate + D,L-lactate |
| *Herbaspirillum seropedicae* SmR1 | Gary Stacey, U. Missouri | RCH2_defined_noCarbon |
| *Marinobacter adhaerens* HP15 | DSM 23420 | DinoMM_noCarbon + L-lactate |
| *Phaeobacter inhibens* BS107 | DSM 17395 | DinoMM_noCarbon + L-lactate |
| *Pseudomonas stutzeri* RCH2 | Romy Chakraborty, LBNL | RCH2_defined_noCarbon + glucose |
| *Shewanella oneidensis* MR-1 | ATCC 700550 | ShewMM_noCarbon + D,L-lactate |
| *Sinorhizobium meliloti* 1021 | ATCC 51124 | RCH2_defined_noCarbon + glucose |
| *Sphingomonas koreensis* DSMZ 15582 | DSM 15582 | RCH2_defined_noCarbon + glucose |

We also studied individual transposon mutants of DVU1186 and DVU2938 from *D. vulgaris* Hildenborough (ATCC 29579). These were obtained by using barcoded variants of the mini-Tn*5* transposon delivery vector pRL27 (Oh et al. 2010), which were delivered by conjugation. Transformants were selected on agar plates (1.5 g/L) with the antibiotic G418 (400 μg/ml) and a rich lactate-sulfate medium. Individual colonies were picked into 96 well plates and characterized by arbitrary PCR and Sanger sequencing (Oh et al. 2010).

We also studied a deletion mutant of DVU0338 from *D. vulgaris* Hildenborough (strain JW9475). This was constructed in a *upp*-background, with JW710 as the parent strain (Keller et al. 2009).

All bacteria were cultured aerobically with shaking, except that *D. vulgaris* Miyazaki (which is strictly anaerobic) was grown without shaking in 18 x 150 mm hungate tubes with a butyl rubber stopper and an aluminum crimp seal (Chemglass Life Sciences, Vineland, NJ) with a culture volume of 10 mL and a headspace of about 15 mL. Media for *D. vulgaris* Miyazaki was prepared in a Coy anaerobic chamber with an atmosphere of about 2% H_2_, 5% CO_2_, and 93% N_2_. Wild type and mutant strains of *D. vulgaris* Hildenborough were grown in a similar way as *D. vulgaris* Miyazaki, but with these differences: the culture volume was only 5 mL; media was degassed for 5 min with 100% nitrogen prior to being autoclaved; and 1.4 mM thioglycolate was used as the reductant in the medium (instead of 1 mM sulfide).

Complementation assays were performed using deletion strains of *thrB, serA,* or *serB* from the Keio collection (Baba et al. 2006). Genes of interest were cloned under the control of the *bla* promoter from pUC19 into the vector pBBR1-MCS5 (Kovach et al. 1995). After sequence verification, we introduced the complementation plasmids into the corresponding knockout strains by transformation. The growth of these strains was tested on M9 minimal media agar plates.

### Mutagenesis of Herbaspirillum seropedicae SmR1

The mutant library of *H. seropedicae* will be described in more detail elsewhere. It was generated using an *E. coli* conjugation donor that contains the plasmid pKMW7 and is an auxotroph for diaminopimelate (Wetmore et al. 2015). The transposon insertion sites were amplified as described previously (Wetmore et al. 2015) and sequenced using Illumina HiSeq2500 in rapid run mode. We identified insertions (supported by at least two reads) at 82,441 different locations in the 5.5 MB genome (NC_014323). We associated 98,021 different barcodes with insertions in the genome (with at least 10 reads for each of these barcodes) and estimated fitness values for 3,878 of the 4,243 non-essential proteins.

### Identification of essential proteins

We identified essential proteins in *H. seropedicae* as described previously for the other bacteria (Price et al. 2016). This approach was validated by (Price et al. 2016; Rubin et al. 2015). Briefly, we limited the analysis to protein-coding genes that were long enough such that the absence of a transposon insertion in the central 10-90% of the gene would be surprising. Protein-coding genes that were long enough (at least 400 nt for *H. seropedicae*) were considered essential if the density of insertion locations (which was normalized by GC content) and the total reads (summed across all insertions) divided by the gene’s length were both less than 20% of the typical protein’s value. Using these thresholds, we identified 472 essential proteins (Supplementary Table 2). These proteins might not be entirely essential but they should be required for good growth in LB.

We also manually classified three steps in biosynthetic pathways as being essential. These involved genes that were not considered in the automated analysis but lack any insertions. In *A. brasilense*, shikimate kinase (AZOBR_RS03225) was originally annotated as a pseudogene, so it was not considered during the automated analysis, but it appears to be essential. Also in *A. brasilense*, aspartyl/glutamyl-tRNA amidotransferase contains three subunits, of which two were automatically identified as essential and one (AZOBR_RS20640) was too short for the automated approach but had no insertions. We classified this step as being essential. Similarly, we classified this step as essential in *P. stutzeri* despite the short length of Psest_3328, which also has no insertions.

### Mutant fitness assays

Most of the mutant fitness assays that we analyzed for this study were described previously (Wetmore et al. 2015; Price et al. 2016). The compendium of mutant fitness assays for *H. seropedicae* will be described elsewhere: it includes growth in minimal media with 26 different carbon sources; growth in minimal media with 1 alternative nitrogen source; and growth in rich media with 12 different inhibitory compounds added. All of this data is available from figshare (https://doi.org/10.6084/m9.figshare.5134837.v1) or the Fitness Browser (http://fit.genomics.lbl.gov).

For this study, we conducted additional mutant fitness assays with amino acids as additional nutrients. These assays were performed and analyzed as described previously (Price et al. 2016). Briefly, a pool of transposon mutants is recovered from the freezer in rich media and grown until it reaches log phase. It is then inoculated at OD_600_ = 0.02 into 5 mL of media in a glass tube and allowed to reach saturation. Gene fitness values are computed by comparing the sample after growth to the sample before growth (i.e., at the time of transfer) via genomic DNA extraction, PCR amplification of barcodes, and sequencing on Illumina HiSeq. For each fitness experiment, metrics of internal consistency and biological consistency were computed and experiments with low quality scores were discarded, as described previously (Wetmore et al. 2015).

Some of the samples were sequenced with a staggered “BarSeq2” primer rather than the primer we used previously. The staggered primer contains 2-5 random nucleotides just downstream of the Illumina adapter. This increases the diversity of the sequence and allows BarSeq to be conducted with the HiSeq 4000.

For the mutant libraries that we published previously (Wetmore et al. 2015; Price et al. 2016), we used the same strains to estimate gene fitness, so that gene fitness values would match for the previously-published results.

### Resequencing of hisD in A. brasilense Sp245

We amplified the *hisD* region from *A. brasilense* by PCR using the primers TCTCCCAGGAGGAGGTGGAC and ATCGCCTTCACGCTGTCCGCATCG. The same primers were used for Sanger sequencing.

### Sequence analysis

To assign genes to TIGRFams (Haft et al. 2013), we used HMMer 3.1b1 44 (Eddy 2011) and the trusted score cutoff for each family in TIGRFam 15.0. TIGRFam assigns enzyme commision numbers to some of its families.

To assign genes to enzyme commission numbers via KEGG (Kanehisa and Goto 2000), we downloaded the last public release of KEGG (from 2011) and we searched for a best hit with over 30% identity and above 80% coverage using RapSearch v2.22 (Zhao et al. 2012). If the best hit was assigned an enzyme commission number by KEGG, then we transferred that annotation to the gene.

To assign genes to enzyme commision numbers via SEED and RAST (Overbeek et al. 2014), we used the SEED server, based on code from http://servers.nmpdr.org/sapling/server.cgi?code=server_paper_example6.pl

These results were viewed using the Fitness Browser (http://fit.genomics.lbl.gov/).

To compute the phylogenetic profile of DUF1638 and the RamA-like protein, we used MicrobesOnline (Dehal et al. 2010). For DUF1638, we used the presence or absence of PF07796. For the RamA-like protein, we used the presence of an ortholog of the RamA-like protein (VIMSS 5050244). However in *Roseobacter sp.* SK209-2-6, the RamA-like protein is split into two proteins (RSK20926_19262 and RSK20926_19267), and we manually classified the RamA-like protein as present in this bacterium.

### Data availability

Mutant fitness data and Sanger sequencing data are available for download at http://genomics.lbl.gov/supplemental/auxo along with supplementary tables 1 and 2. The fitness data can also be viewed in the Fitness Browser (http://fit.genomics.lbl.gov/). The Sanger sequencing data has also been submitted to Genbank (accession KY549926).

### Source code

Code for analyzing fitness data and for the Fitness Browser is available at https://bitbucket.org/berkeleylab/feba

## Appendix 1: Testing the auxotroph predictions of D’Souza and colleagues

(D’Souza et al. 2014) predicted that most bacteria cannot make all 20 amino acids. For example, they predicted that over 30% of sequenced bacterial genomes do not encode the capability to synthesize phenylalanine. Dr. Christian Kost kindly provided us with the list of predictions for phenylalanine auxotrophy (personal communication, September 16, 2014) and we tested these predictions in two ways.

First, of the 10 bacteria we focused on here, they had predictions for five. Two of these five bacteria were predicted to be auxotrophic for phenylalanine (*S. meliloti* 1021 and *D. vulgaris* Miyazaki F). We believe that these predictions are based on the genome sequences of the exact same strains that we studied. This suggests that many of their auxotroph predictions are false positives, but five bacteria is a small sample.

To test the predictions on a larger sample of bacteria, we considered nitrogen-fixing bacteria. These bacteria should be particularly unlikely to be auxotrophic for amino acids, as amino acids provide a more biochemically convenient source of nitrogen than nitrogen gas. Also, nitrogen fixation is usually studied by growing bacteria in the absence of amino acids. So, we compared a list of known nitrogen-fixing bacteria with sequenced genomes (Dos Santos et al. 2012) to the predictions provided by Dr. Kost. We were careful to make sure that the strain identifiers matched. We excluded cyanobacterium UCYN-A from consideration because it is an endosymbiont (Hagino et al. 2013). This left us with predictions for 39 nitrogen-fixing strains. 11 of the 39 strains (28%) are predicted to be auxotrophic for phenylalanine, which is about the same rate as for other bacteria (36%). The difference between the two proportions is not significantly different (P = 0.39, Fisher exact test).

We then examined a random sample of five of the 11 “auxotrophic” nitrogen-fixing bacteria to check that they grow in minimal media. One of these bacteria was *S. meliloti* 1021, which grows in minimal media as long as cobalt and biotin are provided (Watson et al. 2001). For the other four, we found published reports of growth in minimal media with no amino acids. In general, these media contain inorganic salts, buffer, trace vitamins, and a carbon and/or nitrogen source, and none of the media contain amino acids. Specifically, *Clostridium beijerinckii* NCIMB 8052 grows in MP2 medium with acetate as carbon source (Chen and Blaschek 1999). *Clostridium kluyveri* DSM 555 is also known as ATCC 8527 and grows in Stadtman-Burton medium with ammonia or N_2_ as the sole source of nitrogen (Kanamori et al. 1989). *Methylocella silvestris* BL2 grows in NMS media with methane as the sole source of fixed carbon (Theisen et al. 2005). And the cyanobacterium *Trichodesmium erythraeum* IMS101 grows in YBC-II medium with CO_2_ and N_2_ as the major nutrients (Shi et al. 2007).

Overall, we found that the D’Souza and colleagues predicted phenylalanine auxotrophy for many of the bacteria that grow in minimal media. Furthermore, the rate of “auxotrophies” was about the same for bacteria that are verified to grow in minimal media as for other bacteria. This suggests that most of their predicted auxotrophies are spurious.

## Appendix 2: Tests of growth without added amino acids or vitamins

To rule out growth due to the small amounts of nutrients that remain after transfer from rich media into minimal media, we performed multi-transfer growth experiments for seven bacteria (*B. phytofirmans*, *D. vulgaris* Miyazaki F, *H. seropedicae*, *M. adhaerens, P. inhibens*, *P. stutzeri,* and *S. meliloti*). After recovering each of these strains from the freezer in rich media, we transferred them into minimal media (with vitamins but no amino acids), grew them up, and transferred them 1-2 additional times. We observed robust growth of all strains after the final transfer into fresh minimal media. (For this test of *D. vulgaris*, we used 40 mM L-lactate and 20 mM sulfate rather than the usual 60 mM D,L-lactate and 30 mM sulfate.) Also, although we did not test *A. brasilense* Sp245 ourselves, it is reported to grow with nitrogen gas as its sole source of nitrogen (Wisniewski-Dyé et al. 2011), which implies that it can synthesize all 20 amino acids.

We also tested growth of the seven bacteria in defined media with the vitamins omitted. Except for *S. meliloti*, they all grew without added vitamins. *S. meliloti* 1021 is reported to grow in minimal media (with no amino acids) if biotin and cobalt are provided (Watson et al. 2001). The other six strains appear to be able to make all of the necessary vitamins. But it is difficult to be certain that none of our ingredients or glassware was contaminated with small amounts of vitamins.

To test the possibility that our media was contaminated with low but sufficient concentrations of vitamins, we collected mutant fitness data for *B. phytofirmans, H. seropedicae, M. adhaerens, P. inhibens,* and *P. stutzeri* growing in defined media without any added vitamins. We identified auxotrophic phenotypes in more than one organism for mutants in the biosynthesis of NAD, thiamine, and biotin (for NAD biosynthesis genes, in *B. phytofirmans*, *P. inhibens*, and *P. stutzeri*; for thiamine biosynthesis genes, in *H. seropedicae* and *P. stutzeri*; and for biotin biosynthesis genes, in *B. phytofirmans, H. seropedicae, P. inhibens,* and *P. stutzeri).* This suggests that our media did not contain these vitamins at a concentration that was sufficient for efficient growth. Also, in *P. stutzeri*, which contains both a vitamin B12-dependent methionine synthase and a B12-independent methionine synthase, the B12-independent isozyme was important for fitness in the absence of vitamins (gene fitness under -3) but not in the presence of our vitamin mix, which included cyanocobalamin (gene fitness = -0.1). This suggests that our media did not contain sufficient vitamin B12.

Our vitamin mixes also contained vitamin B6 (as pyridoxine), lipoic acid, riboflavin, folic acid, and pantothenate. For all five of these bacteria, the biosynthetic pathways for these vitamins appear to be essential for growth in rich media that contains yeast extract, which explains why we failed to identify auxotrophic mutants in these pathways. These bacteria may not be able to take up these vitamins.

## Appendix 3: Clear candidates whose mutants were not consistently auxotrophic

Most of the clear candidates (that is, genes that were identified by at least two out of three annotation resources as filling gaps in the IMG predictions) were either important for fitness in minimal media, or were likely essential in rich media, or the lack of an auxotrophic phenotype could be explained by genetic redundancy. Here we describe the two exceptions.

First, Psest_1986 from *P. stutzeri* contains a prephenate dehydrogenase domain (PF02153) fused to a 3-phosphoshikimate 1-carboxyvinyltransferase (TIGR01356). Although this gene is expected to be required for tyrosine biosynthesis, it was not important for fitness in most defined media conditions. The gene’s mutant fitness was estimated from just two mutant strains, both of which lie in the linker region between the two domains. The location of these strains is striking because the linker region contains just 40 of the 747 amino acids of Psest_1986. Our interpretation is that both enzymatic activities are essential, and that these insertions were recovered because they allowed both domains to be expressed as separate proteins.

Second, DvMF_1902 from *D. vulgaris* contains a D-isomer specific 2-hydroxyacid dehydrogenase catalytic domain (PF00389) and is annotated as D-3-phosphoglycerate dehydrogenase by both KEGG and SEED. It is 39% identical to a characterized D-3-phosphoglycerate dehydrogenase (MMP1588; (Helgadóttir et al. 2007)). This activity is expected to be required for serine biosynthesis, as a related strain of *Desulfovibrio* appears to use the standard pathway of serine synthesis via phosphorylated intermediates (Germano and Anderson 1969). Also, the isotope labeling of serine in another strain of *D. vulgaris* is consistent with the standard pathway (Tang et al. 2007). Nevertheless, DvMF_1902 was not important for fitness in defined media (all fitness values were -0.5 or higher). Although we did not identify any other strong candidate genes for this activity, DvMF_1902 could be redundant with another dehydrogenase.

## Acknowledgements

This material by ENIGMA-Ecosystems and Networks Integrated with Genes and Molecular Assemblies (http://enigma.lbl.gov), a Scientific Focus Area Program at Lawrence Berkeley National Laboratory is based upon work supported by the U.S. Department of Energy, Office of Science, Office of Biological & Environmental Research under contract number DE-AC02-05CH11231.

